# Individual differences in the engagement of habitual control over alcohol seeking predicts the development of compulsive alcohol seeking and drinking

**DOI:** 10.1101/2020.10.08.331843

**Authors:** Chiara Giuliano, Mickaël Puaud, Rudolf N. Cardinal, David Belin, Barry J. Everitt

## Abstract

Excessive drinking is an important behavioural characteristic of alcohol addiction, but not the only one. Individuals addicted to alcohol crave alcoholic beverages, spend time seeking alcohol despite negative consequences, and eventually drink to intoxication. With prolonged use, control over alcohol seeking devolves to anterior dorsolateral striatum, dopamine-dependent mechanisms implicated in habit learning and individuals in whom alcohol-seeking relies more on these mechanisms are more likely to persist in seeking alcohol despite the risk of punishment. Here, we tested the hypothesis that the development of habitual alcohol-seeking predicts the development of compulsive seeking and that, once developed, it is associated with compulsive alcohol drinking. Male alcohol-preferring rats were pre-exposed intermittently to a two-bottle choice procedure, and trained on a seeking–taking chained schedule of alcohol reinforcement until some individuals developed punishment-resistant seeking behaviour. The associative basis of their seeking responses was probed with an outcome-devaluation procedure, early or late in training. After seeking behaviour was well established, subjects that had developed greater resistance to outcome-devaluation (were more habitual) were more likely to show punishment-resistant (compulsive) alcohol seeking. These individuals also drank more alcohol, despite quinine adulteration, even though having similar alcohol preference and intake before and during instrumental training. They were also less sensitive to changes in the contingency between seeking responses and alcohol outcome, providing further evidence of recruitment of the habit system. We therefore provide direct behavioural evidence that compulsive alcohol seeking emerges alongside compulsive drinking in individuals that have preferentially engaged the habit system.

## Introduction

Individuals with severe alcohol use disorder (AUD) have impaired control over alcohol drinking, but they also spend considerable amounts of time and effort seeking and obtaining alcohol. Although these two diagnostic characteristics of AUD are related^1,2^, they are regulated by distinct neural and psychological processes^3–5^.

Chronic alcohol drinking, especially through intermittent access resulting in escalated intake^6,7^, leads to neurotransmitter, plasticity, and structural changes in the anterior dorsal lateral striatum (aDLS)^8^, such as increased glutamate release and decreased GABA-mediated inhibition at medium spiny neuron synapses, with associated alterations in LTP and LTD^9,10^. In primates, chronic alcohol drinking interspersed with periods of abstinence results in increased dendritic spine density and enhanced glutamatergic transmission in the putamen (analogous to the aDLS in rodent brain), but not in the caudate nucleus, while GABAergic transmission is selectively suppressed in the putamen of monkeys who drink the greatest amounts of alcohol^11^. Furthermore, the emergence of compulsive alcohol seeking^12^ in rats has been shown to be predicted by reliance on, and an inability to disengage, dopamine-dependent mechanisms in the aDLS^13^.

These alterations by alcohol of aDLS function^14,15^ and its emergent control over alcohol seeking has been linked to a transition from goal-directed to habitual drug seeking^14,16^ as shown by the development of resistance to the devaluation of alcohol by lithium chloride aversion or sensory-specific satiety^9,14,15,17,18^. Additionally, habitual responding for alcohol develops more rapidly than for food^19^ or a sucrose reinforcer^14^ and depends on a shift from posterior dorsomedial striatum (pDMS) to aDLS control over responding^14,20^.

While there is a link between the development of habitual and compulsive alcohol seeking^13^, the relationship to compulsive drinking is less clear. In particular, it is uncertain whether increased alcohol consumption causes the development of aDLS-dependent seeking habits and compulsion, or develops in parallel with (or is a consequence of) these behavioural transitions.

In the present experiments, we used our established seeking–taking chained schedule of alcohol reinforcement, which also supports the probabilistic punishment of seeking responses^12,13^. We investigated the action–outcome (A–O) versus stimulus–response (S–R) associative structure underlying alcohol seeking, at different time points during a long history of alcohol use, in alcohol-preferring (P) rats^21–23^. We also assessed the development of compulsive (quinine-resistant)^24–28^ alcohol drinking.

In the seeking–taking chained schedule^29,30^, “seeking” responses are spatially and temporally distinct from “taking” responses. An animal can only gain access to a taking lever, and then the opportunity to drink alcohol, by pressing a seeking lever, but seeking responses are never directly associated with alcohol. We devalued the ultimate outcome of the seeking behaviour by extinguishing the taking link of the chain (via daily sessions of responding on the taking lever alone without alcohol delivery) according to our established procedures^16,31^ and performed this manipulation at time points previously shown either to engage, or not engage, DLS dopamine-dependent mechanisms (short vs long training)^13^.

Compulsive alcohol seeking was assessed by punishing, unpredictably, the completion of some seeking response cycles (instead of presenting the taking lever), so that animals had to risk punishment in order to take alcohol^12,13,32^. Punishment was never associated with taking responses or the delivery of alcohol. We were therefore able to test the hypothesis that animals that were insensitive to reinforcer devaluation, and were responding habitually, were more likely to develop compulsive alcohol seeking. We further tested this hypothesis by investigating whether compulsive and non-compulsive rats were differentially sensitive to degradation of the contingency between seeking responses and outcome (alcohol delivery), a further test of the S–R nature and emergent inflexibility of alcohol-seeking behaviour^33,34^. Finally, we investigated whether the development of punishment-resistant, compulsive alcohol seeking was associated with compulsive, quinine-resistant, alcohol drinking^28,35–38^.

## Methods and Materials

### Subjects

Male alcohol-preferring (P) rats **(see SOM for details)** obtained from Indiana University Medical Center (Indiana, USA) were group-housed during two weeks of habituation to the animal facility, and then single-housed under a reversed 12h light/dark chain (lights off at 07:00) with food and water always available *ad libitum*. Experiments were performed every other day between 08:30 and 16:00 and were conducted in accordance with the UK (1986) Animal (Scientific Procedures) Act (Project Licence PA9FBFA9F).

### Drugs

Ethanol (EtOH) solutions were prepared as described previously^4^ and **detailed in the SOM**.

### Apparatus

Behavioural training was conducted in 12 operant chambers (Med Associates, St. Albans, VT, USA) as previously described^12^. Lever presses, light stimulus presentation, reward delivery, and data collection were controlled by a computer running Whisker control software^39^.

### Procedures

The series of experiments conducted in this study is summarised schematically in **Figure 1 and detailed in SOM**.

**Figure 1:**
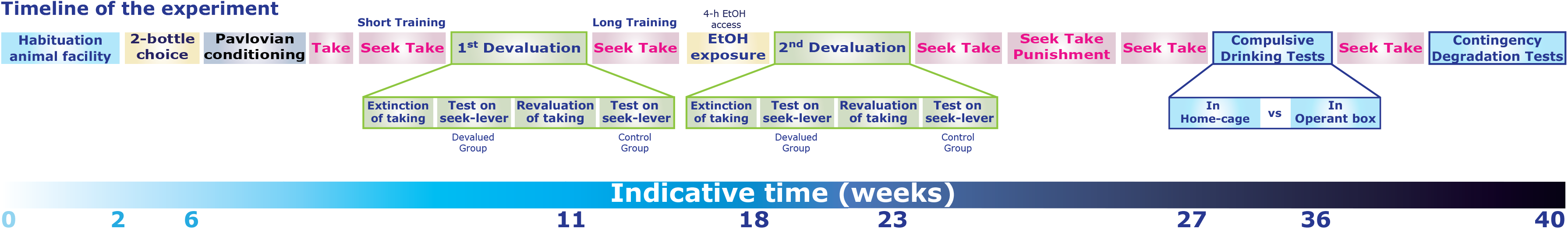
Timeline of the experiments. Alcohol-preferring (P) rats (n=25) were trained in the following stages: **(i) Pavlovian conditioning**, in which rats acquired a light–alcohol association. **(ii) Taking:** rats learned to press a “taking” lever for 15% EtOH under a fixed-ratio-1 schedule. **(iii) Seeking–taking:** rats learned to press a second “seeking” lever to gain access to the taking lever, via a random interval schedule whose parameter increased from 5 to 60 s. Seeking responses were never directly reinforced with alcohol. **(iv) EtOH exposure:** rats were given 4h free access to EtOH in the home cage. **(v) Seeking–taking–punishment:** some seeking cycles were terminated randomly by unpredictable mild foot-shock, rather than insertion of the taking lever. Punishment was never associated with “taking” responses or alcohol delivery. Following this training, rats were assigned to compulsivity subgroups. **(vi) Outcome devaluation:** at two time points (either after three sessions of seeking–taking training, termed **Short Training** or **ST**, or at completion of the full training, termed **Long Training** or **LT**, sensitivity to outcome devaluation was assessed, after either extinction of the taking response (devaluation procedure) or revaluation (control). **(vii) Alcohol intake** was again measured both in the home cage and in the operant chamber, **see SOM**. **(viii) Contingency degradation:** finally, the same subjects underwent sessions in which the contingency of the seeking–taking link was degraded by the non-contingent, free delivery of alcohol.

We confirmed the alcohol-preferring phenotype of P rats in an intermittent two-bottle choice procedure **(see SOM)**. They were then trained instrumentally on a random interval 60s/fixed ratio 1 (RI60/FR1) seeking–taking chained schedule of alcohol reinforcement, as previously described^12,13^ **(see SOM)**.

The development of resistance to outcome devaluation was tested at two time points, in a procedure adapted from Olmstead et al.^40^, Giuliano et al.^12,13^, and Zapata et al.^16^, illustrated in **Figure 1 and SOM**. Critically, the sensitivity of instrumental seeking behaviour to extinction of the taking link was assessed in the absence of alcohol and the taking lever, following several sessions of extinction of the taking lever. This test was conducted after short or long training under the seeking–taking task.

At this point, subjects were identified as compulsive and non-compulsive according to their persistent seeking responses despite the risk of punishment, quantified as the number of completed cycles over the last three days of exposure to 0.45 mA footshocks delivered randomly on completion of some seeking cycles^13^.

After two additional “baseline” sessions of the seeking–taking chained schedule under RI60/FR1, compulsive drinking behaviour was tested as the persistence of alcohol seeking or drinking despite adulteration with bitter-tasting quinine^35^ **(see SOM)**.

After four further re-baseline sessions of the seeking-taking chained schedule under RI60FR1 following the compulsive-drinking test, the sensitivity of alcohol seeking to contingency degradation was investigated, to establish the associative nature of instrumental responding, as **detailed in the SOM**.

### Data and Statistical analyses

Data are presented as means ± SEM, individual data points, or box plots (quartile boxes with minimum/maximum as whiskers). Analyses, **detailed in the SOM**, were carried out across the whole group (dimensional analyses) and in subpopulations (via analyses of variance, ANOVAs) using SPSS 25 (IBM, USA).

Two-tailed values of p < .05 were considered statistically significant. Significant ANOVA main effects and interactions were analysed further using Sidak’s post-hoc test or Dunnett’s test (when comparing multiple time points to a single baseline) as appropriate. Effect sizes are reported as partial eta squared (η_p_^2^)^41^.

## Results

The alcohol-preferring phenotype of the 25 P rats that completed the experiment was confirmed over twelve two-bottle choice sessions under intermittent access^7,42^ **(Figure S1, see SOM for more details)**. Rats subsequently shown to develop alcohol-seeking habits acquiring high drinking levels slightly earlier than their counterparts **(Figure S1, see SOM for more details)**. Rats were then trained to self-administer alcohol over four sessions under continuous reinforcement, by the end of which they had all reached the maximum number of rewards per session available (45 deliveries of 0.1 mL 15% EtOH) **(Figure S2A, see SOM for more details)**. Rats were introduced to a RI5/FR1 seeking–taking schedule of alcohol reinforcement for three sessions (**Short Training; ST**), eventually reaching 113±15.89 seeking and 35±4.22 taking lever presses by the final 2-hour session **(Figure S2B)**.

The goal-directed or habitual nature of early alcohol seeking was tested by measuring the sensitivity of alcohol-seeking responses to the devaluation of their outcome, namely access to the taking lever, across two tests during which only the seeking lever was presented, either after extinction of the taking-lever-to-alcohol link (devalued condition), or the resumption of alcohol taking and revaluation of the link (revalued, control condition)^16^.

Withholding alcohol delivery resulted in extinction of “taking” responses across 17 sessions [time: F_16,384_=86.01, p<.001, η_p_^2^=.78]. Rats made on average 9±.97 lever presses over the last two sessions of extinction, a reduction of about 90% compared to the first day of extinction **(Figure S2C-D, see SOM for more details)**. Rats subsequently identified as being compulsive showed a higher initial level of responding (less initial extinction) than non-compulsive rats, but eventually showed the same degree of extinction **(Figure S4, see SOM for more details)**. Drug-taking responses returned to pre-extinction levels when alcohol access was resumed (under continuous reinforcement, for 2 sessions).

Across the whole group, alcohol seeking after short training was sensitive to outcome devaluation. Extinction of the taking link resulted in a marked reduction, of around 50%, in responses on the seeking lever (t=5.37, df=24, p<.001) **(Figure 2A)**.

**Figure 2:**
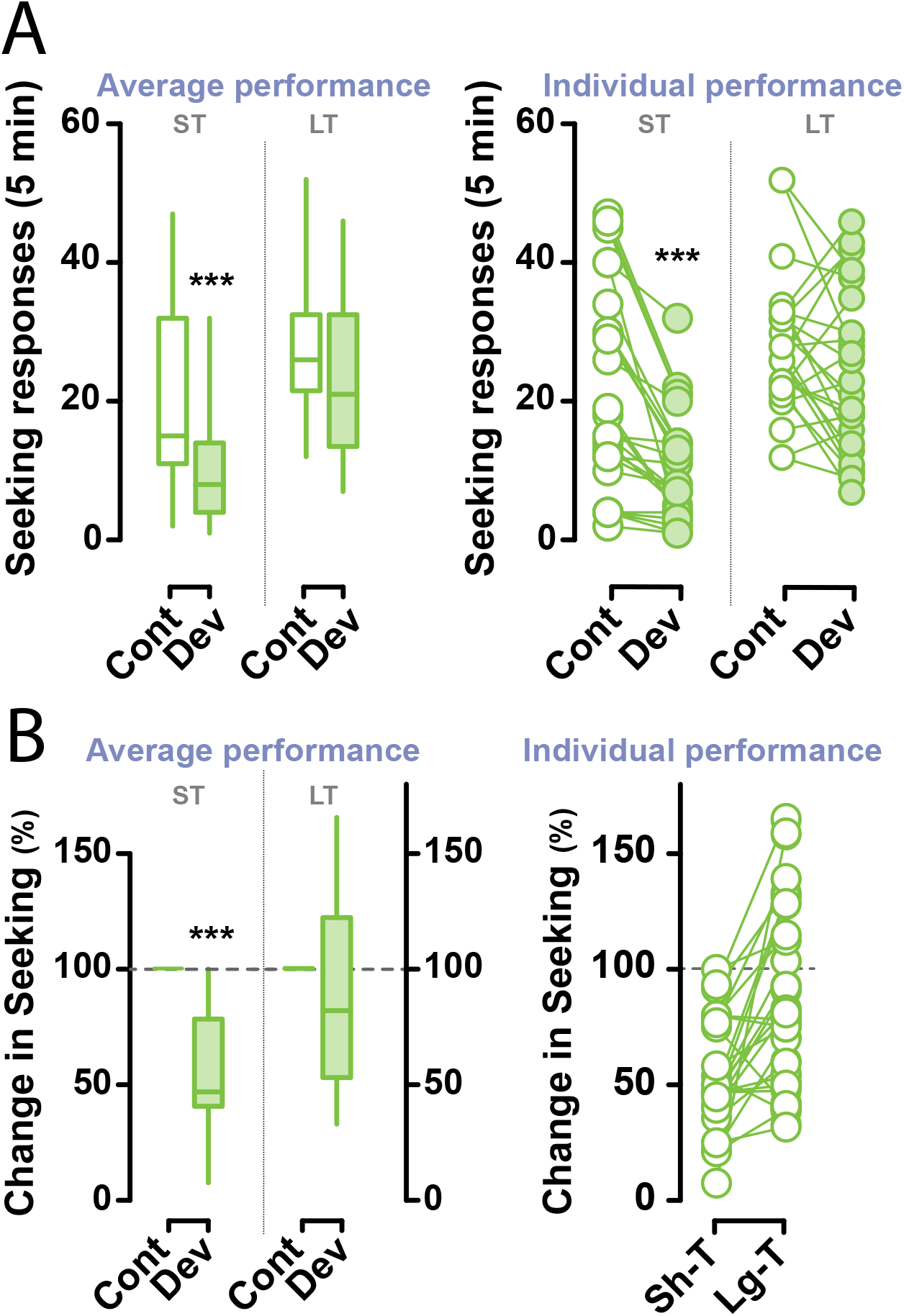
Devaluation testing after short and long training. After short or long training on the alcohol seeking–taking schedule, responses on the drug seeking lever only were measured during 5 min tests after extinction of the taking link (devalued condition) and after revaluation of the alcohol taking link (control condition). **A)** Total seeking responses per session (left, mean; right, per subject). **B)** The magnitude of the devaluation effect was calculated as the number of seeking responses after devaluation, as a percentage of seeking responses under the revalued condition (left, mean; right, per subject; *** p< .001 versus revalued condition).

Following this first test, rats were given more extensive experience in the reinforced seeking– taking task under RI60/FR1 (**Long Training, LT**). They eventually reached 320±35.07 seeking and 24±.34 taking lever presses over the last two sessions of training (in which the number of cycles was limited to 25) **(Figure S2E, see SOM for more details)**.

Withholding alcohol delivery resulted in similar levels of extinction of the taking response, albeit over 24 sessions, to that seen in the earlier performance test [time: F_23,552_=91.82, p<.001, η_p_^2^=.79]. Rats later identified as compulsive again showed an initial higher level of responding in extinction, but subsequently extinguished to the same degree as their non-compulsive counterparts **(Figure S4, see SOM for more details)**. All rats eventually made on average 10±.98 lever presses over the last two extinction sessions, a reduction of 90% from the first session **(Figure S2F-G, see SOM for more details)**. Drug-taking responses returned to pre-extinction levels when access to alcohol was reinstated (under continuous reinforcement for 2 sessions, data not shown). However, across the whole group, this devaluation of the taking link no longer had an effect on seeking (t =−1.93, df=24, p=.07). Analysis of the number of seeking lever presses during test sessions revealed a significant interaction between training experience and devaluation [F_1,24_=4.73, p=.040, η_p_^2^=.17] **(Figure 2A)**.

Analysis of performances as percentage change in alcohol seeking, shown in **Figure 2B**,[training: F_1,24_=18.41, p<.001, pη^2^=.43], revealed that not all individuals were equally sensitive to outcome devaluation at the two time points: 32% of the population remained sensitive to outcome devaluation even after long training **(Figure 2B)**. Rats were stratified according to their sensitivity to devaluation at the long training time point. They were classified as devaluation-sensitive (DS, n=8) if they decreased their seeking responses after outcome devaluation by 40% or more (on average 45.20% ±3.44), or devaluation-resistant (DR, n=17) if their seeking decreased by less than 40% (on average 107.87% ±7.42). These identified sub-groups [group: F_1,23_=31.59, p<.001, η_p_^2^=.58], confirmed by a *k*-means cluster analysis, showed a very different trajectory with regards to their sensitivity to the devaluation of the seeking link [training × group × devaluation: F_1,23_=11.92, p=.002, η_p_^2^=.34]. Post hoc analyses confirmed that while DR and DS rats did not differ from each other after short training, they did in the devalued (p=.003), but not control condition, after long training **(Figure 3A)**. Similarly, when performance was compared between devalued and control conditions for the DS and DR rats independently, the former showed a decrease in responding in the devalued condition after both short- and long-training (p=.009 and p<.001, respectively) whereas the latter (DR) showed this decrease only after short training (p=.009) **(Figure 3A)**.

**Figure 3:**
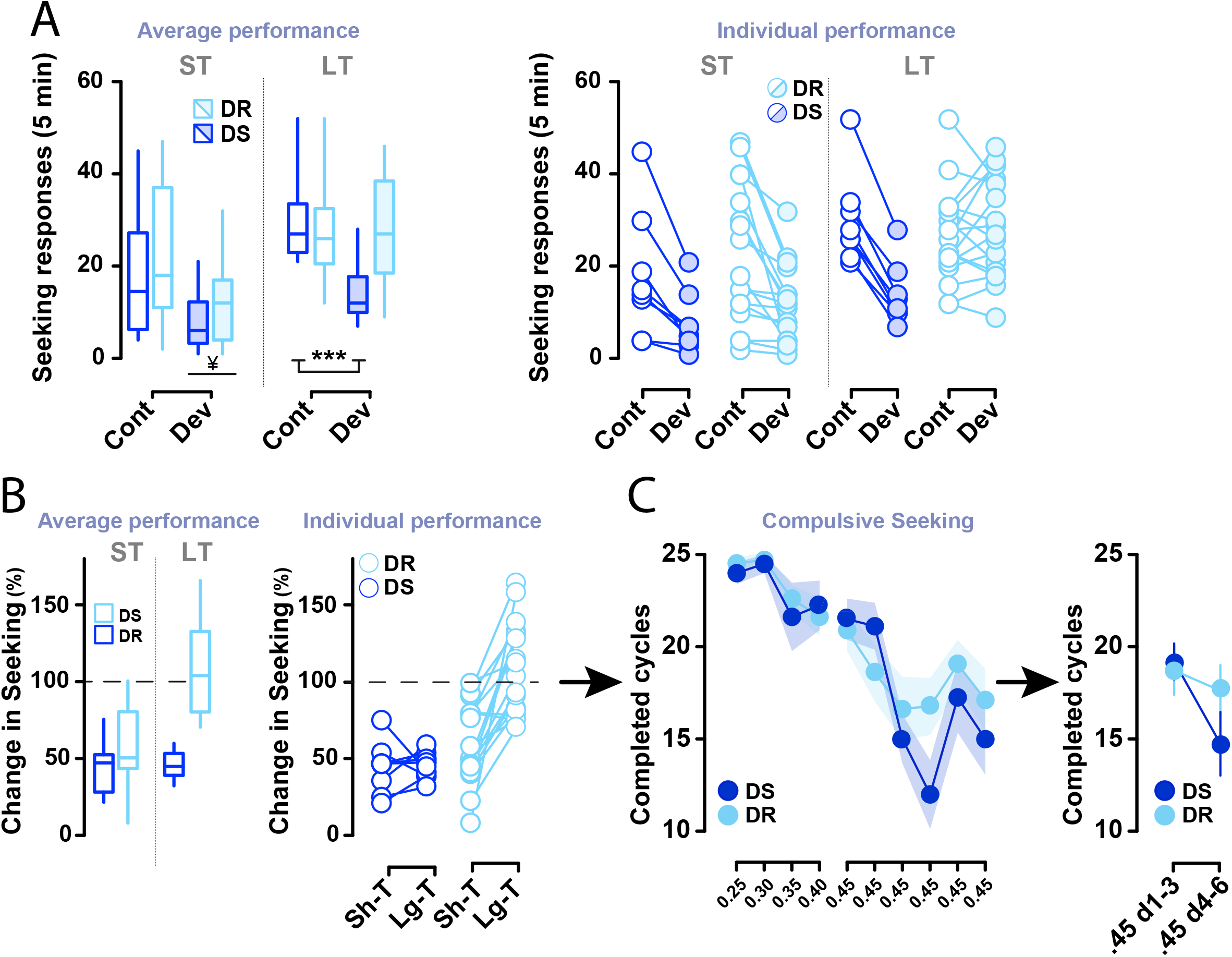
Resistance to outcome devaluation after long training predicts compulsivity. According to the magnitude of the devaluation effect after long training, subjects were assigned to a devaluation-sensitive (DS, blue, n=8) or devaluation-resistant (DR, light blue, n=17) group. **A)** Total seeking responses per 5-min devaluation session (left, mean; right, per subject) for the DS and DR groups. **B)** Magnitude of the devaluation effect, as for **Figure 2**, for DS and DR groups. **C)** Subjects showing higher resistance to devaluation after long training also showed higher resistance to punishment. They completed more seeking–taking cycles when seeking responses were randomly punished by a 0.45 mA, 0.5 sec foot shock. ¥ p< .01 devalued vs control condition; *** p< .001 devalued vs control condition in devaluation-sensitive rats.

The development of alcohol seeking that is resistant to devaluation is a behavioral expression of a shift from goal-directed action (sensitive to outcome devaluation) to habitual alcohol seeking (resistant to devaluation). We hypothesized that the individual variability in the development of *habitual* alcohol seeking would predict the tendency to develop *compulsive* alcohol seeking, identified by persistent seeking despite punishment.

The probabilistic punishment of seeking responses decreased them across all subjects, “dose”-dependently related to foot-shock intensity [time: F_9,216_=9.18, p<.001, η_p_^2^=.28]. DS and DR rats showed no differential response to weaker shocks that didn’t decrease whole-group responding (up to 0.45 mA) [time: F_9,207_=22.69, p<.001, pη^2^=.50; time × group: F_9,207_=1.68, NS; group: F_1,23_<1, NS]. However, over the last sessions, at the higher 0.45 mA intensity, DR rats were more resistant than DS rats to the punishment of seeking [time: F_5,115_=13.41, p<.001, η_p_^2^=.37; time × group: F_5,115_=13.41, p=.012, η_p_^2^=.12; group: F_1,23_<1, NS] **(Figure 3C, left panel)**, such that DR continued to make seeking responses when punishment was at that intensity, whereas DS rats decreased their alcohol seeking to a level significantly different from the first three sessions of punishment at 0.45 mA (time × group: F_1,23_=4.84, p=.038, η_p_^2^=.17; p=.002, first 3 sessions vs last 3 sessions for the DS group), **(Figure 3C, right panel)**.

Marked individual differences were also observed in the persistence of alcohol seeking after seeking was punished. We used the number of seeking-taking cycles completed during the last 3 sessions of punishment to classify rats (via *k*-means cluster analysis)^12,13^ as compulsive (C, n=7), intermediate (I, n=8) or non-compulsive (NC, n=10) [group: F_2,22_=84.16, p<.001, η_p_^2^=.88; time × group: F_4,44_=1.77, NS; time: F_2,44_=7.47, p=.002, η_p_^2^=.25; Sidak’s post-hoc comparisons: C vs I and NC, p<.001, in each comparison] **(Figure S5)**. By definition, C rats showed alcohol seeking that was completely resistant to punishment while non-compulsive rats showed a marked decrease in alcohol seeking. Both the incidence and the qualitative nature of these groups were similar to those previously described using a similar procedure^12,13^.

We hypothesised that individuals that eventually seek alcohol compulsively would also show increased alcohol drinking. Therefore, we retrospectively compared the alcohol intake of C and NC rats, prior to the development of compulsive alcohol seeking. During the instrumental initial training period, rats had been given ten sessions of 4h free-access to 15% EtOH in their home cages. Comparison of the average intake during the first vs the last two sessions showed that all rats drank similar volumes of alcohol, whether or not they subsequently went on to develop compulsive alcohol seeking [session: F_1,15_=.21, NS; group: F_1,15_=1.49, NS; session × group: F_1,15_=.12, NS]. At the dimensional level, the tendency to drink alcohol freely, before punishment, did not predict punishment-resistant seeking behavior either (R^2^=.033, ns) **(Figure 4A, left panel)**.

**Figure 4:**
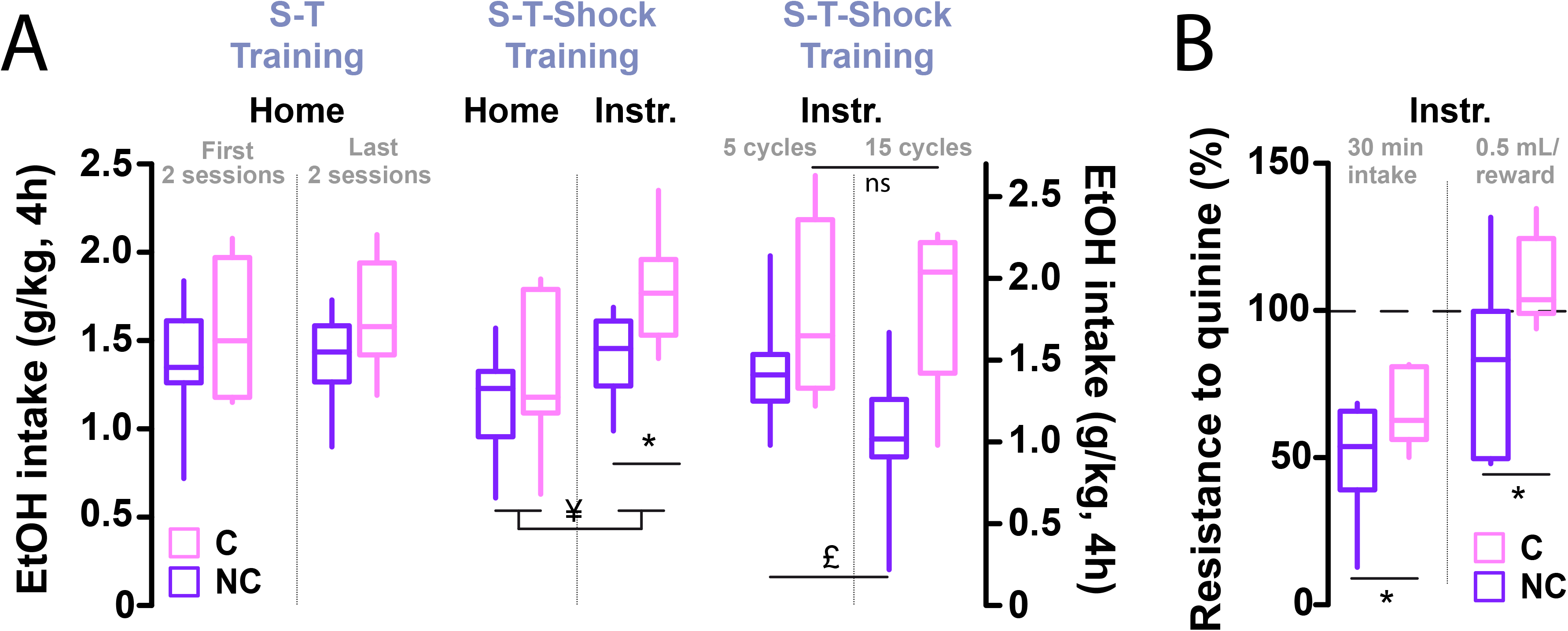
The development of compulsive alcohol seeking is associated with the development of compulsive alcohol drinking. Subjects were assigned to two groups according to the number of completed cycles over the last three days of exposure to 0.45 mA punishment: compulsive (C, pink, n=7) and non-compulsive (NC, purple, n=10). **A)** Drinking behaviour (alcohol intake in g/kg) was assessed in C and NC rats over 4 hour free access challenge sessions (i) over the course of the development of compulsive alcohol seeking (**left panel**), (ii) in the home-cage or the operant chamber after the development of compulsive alcohol seeking (**middle panel**), at time point at which their sensitivity to pre-loading was also assessed following 5 or 15 cycles in the seeking-taking task (iii) (**right panel**). **B)** Resistance to quinine (0.1 g/L) adulteration (expressed as percentage change from baseline consumption of 15% EtOH), an index of compulsive drinking) was assessed (i) over a 30 min of free access period following **(left panel)** or (ii) while performing (**right panel**), 15 seeking-taking chained cycles. ** p< .001, * p< .05 C vs NC. ¥ p< .01 Home vs Instrumental context.

However, following the development of compulsive alcohol seeking in vulnerable rats, C rats drank more alcohol (compared to NC rats) when given the same free access to 15% EtOH in their home cage [group: F_1,17_=7.66, p =.014, η_p_^2^=.34] **(Figure 4A, middle panel: Home)**. C rats also escalated their intake when alcohol was freely available for 4 hours in the instrumental context (Instr.) [group: F_1,15_=5.26, p=.037, pη^2^=.26; group × context: F_1,15_=1.14, ns, pη^2^=.07] (**Figure 4A, middle panel: Instr.)**, where the tendency to drink alcohol was higher than that shown by the population in the home cage [context: F_1,15_=12.69, p=.003, pη^2^=.46]. These observations thereby suggest a loss of control over intake in rats identified as compulsively seeking alcohol.

We tested the hypothesis that C rats had lost control over intake, by measuring their ability to titrate their intake of freely available alcohol according to the quantity of alcohol they had ingested in the immediately preceding seeking–taking session. Rats were given 4h of access to 15% EtOH in their instrumental context immediately following either 5 or 15 seeking–taking cycles in one session. C rats drank more than NC rats [group: F_1,14_=9.02, p=.009, pη^2^=.39]. Additionally, C rats showed a loss of a satiety effect, in that they failed to adjust their intake in response to the amount of alcohol earned in the instrumental context, whereas NC rats drank less in the free-alcohol test after 15 cycles than after 5 cycles [cycles x group: F_1,14_=5.80, p=.030, η_p_^2^=.29; C after 5 cycles vs C after 15 cycles, NS; NC after 5 cycles vs NC after 15, p=.006] **(Figure 4A, right panel)**.

Next, we tested whether the loss of control over alcohol intake shown by C rats was associated with persistence of alcohol drinking despite the negative consequence of quinine ingestion, a widely used test of inflexible, or compulsive, alcohol consumption^28,35–38^. Resistance to quinine (0.1 g/L) adulteration, expressed as percentage change in intake of 15% EtOH **(Figure 4B)**, was measured under two different conditions. First, rats had 30 min access to 15% EtOH adulterated with quinine in the operant chamber after completing 15 cycles of the RI60/FR1 seeking–taking schedule (for unadulterated alcohol). Second, rats underwent 15 cycles of seeking–taking under RI60/FR1, but received 0.5 mL quinine-adulterated 15% EtOH on completion of each cycle.

As predicted, C rats were resistant to quinine adulteration as compared to NC rats. They drank more adulterated alcohol over a 30 min challenge in the same operant box in which the seeking-taking sessions occurred [resistance to quinine: group: F_1,14_=6.54, p=.023, η_p_^2^=.32 **(Figure 4B, left panel)**]. When subjects earned adulterated alcohol during the seeking–taking chained schedule, C rats drank significantly more EtOH than NC rats when compared to baseline intake [resistance to quinine: group: F_1,16_=6.15, p=.026, pη^2^=.30 **(Figure 4B, right panel)**], further demonstrating resistance to quinine adulteration even within the seeking-taking-drinking chain.

Together these results suggest that compulsive rats both seek alcohol habitually and lose control over alcohol intake, which they maintain when adulterated with quinine, indicating that they no longer monitor the consequences of their behaviour. We therefore tested the hypothesis that C rats would be insensitive to degradation of the seeking response contingency. 24 subjects underwent three sessions of seeking-taking chained schedule in which response non-contingent (free) 0.1 mL alcohol deliveries occurred when the seeking lever was extended, in addition to the 0.1 mL alcohol deliveries that were contingent on taking lever responses. This modification of the schedule enabled an assessment of rats’ sensitivity to the causal relationship between seeking actions and their consequences (i.e. pressing the seeking lever to gain access to the take lever in order to obtain and drink alcohol). Since seeking responses at baseline level were higher in C than NC rats, data are expressed as percentage change from baseline **(Figure 5A-B)** (i.e. performance during seeking-taking chained schedule with 0.1 mL 15% EtOH delivered contingent on taking lever responses).

**Figure 5:**
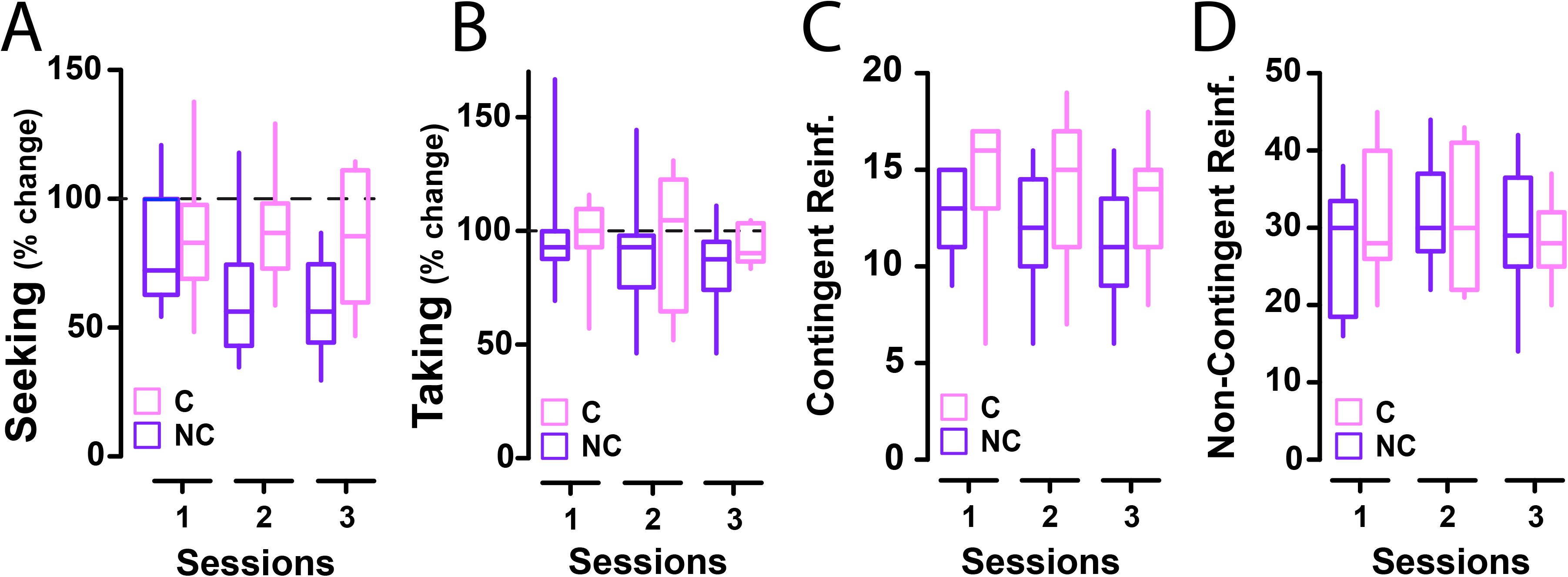
The development of compulsive alcohol seeking is associated with insensitivity to degradation of the instrumental contingency. Compulsive (C, pink, n=7) and non-compulsive (NC, purple, n=9) rats underwent three sessions where the contingency between seeking and taking responses was degraded by non-contingent reward delivery. **A)** Seeking responses, expressed as percentage change from baseline, decreased over time in C rats during the 3 1h-sessions. The contingency degradation procedure did not affect taking responses **(B)**, the number of reinforcers delivered contingent upon the taking lever (FR5 schedule) **(C)**, the number of non-contingent reinforcers delivered when the seeking lever was extended **(D)**, in C and NC rats.

As predicted, C rats maintained significantly higher levels of seeking responses than NC rats during non-contingent alcohol delivery [group: F_1,14_=5.61, p=.033, η_p_^2^=.29; day: F_2,28_=1.51, NS; day × group: F_2,28_=1.40, NS] **(Figure 5A)**. Taking responses **(Figure 5B)**, and the number of contingent **(Figure 5C)** and non-contingent reinforcers **(Figure 5D)**, were not altered [taking: F_1,14_<1, NS; contingent reinforcers: F_1,14_=2.45, NS; non-contingent reinforcers: F_1,14_<1, NS].

## Discussion

We have shown that in rats responding for alcohol in a chained seeking–taking schedule, individual variability in the development of insensitivity to devaluation of the seeking outcome^16,40^ predicted the subsequent development of compulsive, punishment-resistant^4,15^ alcohol seeking. Once established, compulsive alcohol seeking was then shown to be insensitive to degradation of the contingency between seeking and the taking outcome. The development of compulsive alcohol seeking was not preceded or predicted by a higher level of alcohol intake, as we had shown previously^12^. However, it was associated with an escalation of free alcohol intake, especially in the alcohol seeking context, and also insensitivity to adulteration of alcohol with quinine. This is indicative of the emergence of a loss of control over intake and a compulsive drinking phenotype^35,37^. These results parallel the previous demonstration that escalation of cocaine intake results from, but is not causally involved in, the development of compulsive self-administration of the drug^43,44^. These data further suggest that the neurobehavioural basis of alcohol preference or high alcohol intake is dissociable from that of the vulnerability to develop compulsive alcohol seeking and drinking.

Previously, using the same task, we showed that emergence of control over alcohol seeking by dopamine-dependent mechanisms in the aDLS was necessary for the development of compulsive alcohol seeking, and the inability to disengage this aDLS control in the face of punishment further characterised the compulsive state^4^. The present data provide behavioural evidence for the interpretation that aDLS control over seeking behaviour indicates engagement of the habit system^45^. By devaluing the seeking response outcome through extinction of the taking link of the chain^16,40^, habitual seeking was revealed in some individuals after several months of exposure to alcohol, whether through instrumental training or extended alcohol intake in the home cage.

These results are consistent with the demonstration (using a single lever task and sensory-specific satiety to devalue alcohol) that responding for alcohol is goal-directed and dependent on the pDMS after 4 weeks of drinking but becomes resistant to devaluation and dependent on the aDLS after 8 weeks of drinking^14^. This shift from goal-directed to habitual responding over a long reinforcement history under random interval schedules, as well as a shift to control by the aDLS, has also been shown for cocaine ^16,40^ and in several pioneering studies with ingestive food rewards^34,45,46^.

While the resistance to outcome devaluation was evident across all subjects, further analysis revealed clear individual differences in the trajectories of this transition. Two sub-groups were identified, one in which individuals reduced their seeking responses by 40% or more (i.e. were sensitive to outcome devaluation) and another comprising individuals that were resistant to devaluation and maintained their responding. This is consistent with earlier observations of individual variability emerging in the sensitivity of drug seeking responses to inactivation of^16^, or dopamine receptor blockade in^13^, the aDLS.

Devaluation-resistant individuals were more likely subsequently to show punishment-resistant, compulsive alcohol seeking, reflected in a higher number of completed seeking-taking cycles per session when seeking intervals were unpredictably punished. Similarly, retrospective analysis showed that rats who persisted compulsively in seeking alcohol despite the risk of punishment, maintaining their responding at pre-punishment levels, were those that had previously developed devaluation-resistant seeking behaviour.

These differences could not be attributed to different degrees of alcohol preference among the P rat population, or differences in the acquisition of instrumental seeking behaviour. However, rats that eventually revealed themselves to be compulsive consistently showed less early sensitivity to extinction. This was not due to an inability to learn the new response–“no US” association that drives extinction, since compulsive and non-compulsive rats reached the same low level of responding at the end of each extinction challenge. Instead, it suggests compulsive rats were either more motivated for alcohol, as previously established under a progressive ratio schedule^12^, or had a less flexible instrumental response system (despite their sensitivity to devaluation at some points). This observation is concordant with the loss of flexibility demonstrated in compulsive rats in that they cannot disengage aDLS control over alcohol seeking following a change in the seeking environment resulting from the introduction of probabilistic punishment^13^.

The present results lend considerable support to the hypothesis that the engagement of the habit system in rats seeking alcohol, shown by resistance to outcome devaluation (present data) and recruitment of the aDLS^13^, predicts individual vulnerability to develop compulsive seeking, and characterises its phenotype.

We investigated this further by degrading the contingency between seeking responses and outcome, in compulsive and non-compulsive individuals. Contingency degradation is frequently used to test the associative structure underlying instrumental responding, and is typically achieved by the response-independent, unexpected delivery of ‘free’ outcomes^34,47,48^, (in this case, alcohol delivery independent of seeking responses). If seeking is under action– outcome control, it should decrease when free alcohol reinforcers are delivered, but will not decrease if seeking responses are habitual^48^. Compulsive rats maintained significantly higher levels of seeking under contingency degradation conditions, while non-compulsive rats decreased their seeking.

Since the seeking link of the chain was under the control of a random interval schedule, the taking lever was still presented when these intervals elapsed provided animals were still responding, even though compulsive and non-compulsive rats did so at different rates. It was therefore possible to assess whether contingency degradation during the seeking intervals influenced the performance of taking responses under the fixed-ratio-5 schedule component. Compulsive and non-compulsive rats did not differ in their taking responses (or in the number of response-contingent reinforcers received), even though their preceding seeking responses had been differentially affected by the free delivery of alcohol. This further emphasises that seeking and taking responses in chained schedules are under dissociable control, consistent with our previous demonstration that when seeking responses devolve to control by the aDLS, taking responses do not^4^. While there is a strong tendency for seeking responses to shift from goal-directed actions to become habits over time, there are few data to suggest that taking responses do so; they remain instead goal-directed (see^3^ for review). Similarly, in rats responding for multiple food reinforcers, goal-directed and habitual seeking responses have been shown to co-exist, with control over behaviour shifting between goal-directedness and habits on the same day and in the same individual when trained under ratio versus interval schedules of reinforcement, respectively^49^. Insensitivity to reinforcer devaluation, or contingency degradation, appear to affect seeking responses primarily, or more readily, suggesting that in an instrumental chain, responses more distal to the goal are more likely to come under habitual control^18^. However, conditions may exist (for example, a more extended period of training), that eventually result in a loss of goal-directedness in taking responses.

There is consistent evidence that the dorsal striatum of rodents is highly sensitive to the effects of ethanol, and that chronic alcohol intake or intermittent alcohol drinking result in structural, neurochemical and plasticity adaptations in the aDLS, or putamen in primates^8–11,15,50–54^. These data encourage the view that the progressive engagement of the habit system is a consequence of these alcohol-induced adaptations^14,15,55–58^. However, they might also be related to changes in alcohol drinking, as suggested by the finding that long-term alcohol exposure is associated with upregulation of dopamine D3 receptors in the dorsal striatum, but not the ventral striatum, and D3 receptor blockade leads to a reduction in alcohol intake^51^.

All rats preferred alcohol over water equally in a two-bottle choice setting. Even after a prolonged history of instrumental training to respond for alcohol, devaluation-sensitive, devaluation-resistant, compulsive, and non-compulsive rats all drank similar amounts of alcohol. Thus, neither alcohol preference nor the volumes of alcohol drunk predicted the later development of compulsive seeking, confirming our earlier data^4^. However, once compulsive alcohol seeking had emerged, compulsive rats drank more alcohol (when freely available) than non-compulsive rats and this difference was accentuated in the environment in which their compulsivity had developed. This is consistent with studies indicating the important role of the context of drug use in enhancing craving and the performance of ethanol-seeking behaviour^59,60^.

Compulsive (but not non-compulsive) rats were also incapable of adjusting their alcohol consumption in response to the amount of alcohol recently consumed as a reinforcer in the seeking–taking task. This further indicates that their performance was inflexible and not determined by outcome value. Moreover, compulsively seeking (but not non-compulsive) rats persisted in drinking alcohol adulterated with quinine, indicating that their drinking had also developed a compulsive quality, as seen in recent studies in mice^27,38^.

Chronic ethanol exposure has been shown to result in an altered excitatory–inhibitory balance in medium spiny neurons in the aDLS, favouring increased aDLS output that may therefore be associated with both inflexible, compulsive alcohol seeking and also inflexible, compulsive drinking insensitive to changes in taste adulteration as shown here and in other studies^57^. Aversion-resistant alcohol intake has also been shown to be characterised by less variable, more automatic responding, as well as a greater tendency to do so^25^.

The neural mechanisms and circuit basis of these complex changes in alcohol seeking and consumption have yet to be fully determined. In rats, preference for alcohol and the future development of compulsive alcohol seeking^28^ and drinking^24^ have been linked to individual differences in the expression of the GABA transporter GAT3 in the amygdala, while compulsive drinking has been linked (in mice) to altered function in a medial prefrontal cortex–dorsal periaqueductal gray circuit involved in punishment avoidance or resilience^38^. However, the observation that only compulsive P rats develop quinine-resistant, compulsive alcohol drinking suggests that the neural mechanisms underlying the universal tendency of P rats to drink high volumes of alcohol do not necessarily lead to the development of aversion-resistant compulsive drinking, even though P rats tend to drink more quinine-adulterated alcohol than the Wistar rats from which they were originally derived^61^.

Taken together, the present results show that the tendency to develop compulsive alcohol seeking is predicted by the development of habitual alcohol seeking, but not by alcohol preference or alcohol intake. The increased alcohol intake that develops in compulsive seekers is also inflexible, being insensitive to “pre-loading” and resistant to adulteration by quinine. These data suggest that the compulsive nature of alcohol drinking emerges alongside compulsive alcohol seeking habits in vulnerable individuals.

## Supporting information

SOM

## Acknowledgements

The present study was funded by BBRF NARSAD Young Investigator Grant (RG95599 GRANT ID: 27353 to CG) and a Medical Research Council (MRC) Programme Grant (MR/N02530X/1). RNC’s research is funded by the MRC (grant MC_PC_17213 to RNC).

The experimental work was carried out under Home Office Project Licence PA9FBFA9F held by A. Milton. The production of the P rats was funded by the R24 Alcohol Research Resource Award grant (U24AA015512) from NIAAA. The authors would like to thank Zohal Nikzad for technical assistance and Larry Lumeng, Richard Bell and Rebecca Jane Smith at the Indiana University School of Medicine for facilitating the provision of the selected lines of rats.

## Authors contribution

CG, DB and BJE were responsible for the study concept and design; CG performed experiments assisted by MP; CG and DB analysed the data; RNC wrote the software used in the experiments; CG, BD and BJE wrote the manuscript. All authors critically reviewed and edited content and approved the final version for publication.

