## Supplementary material for "Individual differences in the engagement of habitual control over alcohol seeking predicts the development of compulsive alcohol seeking and drinking": SOM

#### TITLE:

#### Abbreviated Title:

Resistance to outcome devaluation predicts compulsivity

#### Authors:

Chiara Giuliano, Mickaël Puaud, Rudolf N. Cardinal, David Belin and Barry J. Everitt

### Supplemental Online Material

#### Methods and Materials

##### Subjects

Male alcohol-preferring (P) rats ~30 days old, n=26, ~250 g at the start of the experiments) obtained from Indiana University Medical Center (Indiana, USA) were group-housed during two weeks of habituation to the animal facility, and then single-housed under a reversed 12h light/dark chain (lights off at 07:00) with food and water always available ad libitum. Experiments were performed every other day between 08:30 and 16:00 and were conducted in accordance with the UK Animal (Scientific Procedures) Act 1986 (Project Licence PA9FBFA9F). Two rats had to be killed due to ill health: 25 rats completed the set of experiments up to the development of compulsive seeking behaviour, 24 rats continued the remaining set of experiments (i.e. quinine adulteration and contingency degradation manipulations).

##### Drugs

Ethanol (EtOH) solutions were prepared by mixing 99.8% EtOH (Sigma, UK) in tap water to obtain 10% EtOH (v/v; 2-bottle choice procedure) or 15% EtOH (v/v; instrumental conditioning).

Quinine (Sigma, UK) was dissolved in 15% EtOH solution to obtain the concentration of 0.1 g/L (w/v)[1-3].

#### Procedures

##### *Alcohol-Water Two-bottle choice*

Animals were presented in their home-cage with intermittent concurrent access to one bottle of 10% EtOH (v/v) and one bottle of water for a total of 12 sessions. The bottles were weighed once a day at the same time, and voluntary consumption of the two liquids was assessed every 24h. The bottle position was changed daily in order to avoid any side preference. The volume consumed in each session was calculated as the weight difference between the start and the end of the sessions, minus 1.5 g and 1 g for spillage for EtOH and water respectively (determined as the volume of fluid lost from bottles in an empty cage). Alcohol intake (g/kg) and the percentage of alcohol preference over the total fluid intake (%) were assessed every 24h.

##### *Training under the seeking–taking and seeking–taking/punishment tasks*

The task enabling the measurement of the development of compulsive alcohol-seeking behaviour[4,5] consisted of the following five phases:

**I. Pavlovian conditioning:** Rats were initially trained to associate presentation of a light stimulus with the opportunity to drink 15% EtOH: a 20-s light CS was illuminated during delivery of 0.1 mL 15% EtOH (v/v) in the receptacle between the two levers. Sessions lasted 45 min and rats were exposed on average to 54 light-stimulus associations.

**II. Taking phase:** Each cycle began with the insertion of the randomly assigned taking lever. Rats were trained to press the taking lever under a fixed-ratio 1 (FR1) schedule of reinforcement, which resulted in 5-s CS illumination, extinction of the house light, and delivery of 0.1 mL 15% EtOH. This phase consisted of 4 sessions, in which rats were limited to a maximum of 45 rewards/2h-session.

**III. Seeking–taking phase:** Each cycle in the seeking–taking chained schedule of reinforcement began with insertion of the seeking lever (at a location distinct from the taking lever that was retracted). Seeking lever presses were never directly reinforced, but instead resulted in the extension of the taking lever and the simultaneous retraction of the seeking lever, under a random interval (RI) schedule. Taking lever responses resulted, via an FR1 schedule, in the illumination of the stimulus light above the taking lever for 5 s, delivery of 0.1 mL of 15% EtOH, and a time-out of 2 min, during which both levers were retracted. Four RI schedules were used across the training period: RI5, RI15, RI30, and RI60, with increments occurring after two consecutive sessions under each RI schedule. Sessions were limited to 2h.

**IV. Free alcohol exposure:** Rats underwent ten sessions with 4h access to a 15% EtOH bottle in the home-cage. Alcohol intake (g/kg) was assessed for 4h.

Interspersed between the extended alcohol sessions (ten in total) were additional sessions of training on the seeking–taking chained schedule under RI60. During the last two sessions under RI60, rats were limited to a maximum of 25 seeking–taking response cycles per 2h session, in order to establish a baseline level before test 2 and the subsequent introduction of the probabilistic punishment contingency, during which the proportion of rewarded and punished trials was kept constant across 25 cycles.

Within phases III and IV, a sub-distinction was made according to the number of sessions performed:

- **Short training:** involved only 3 sessions under RI5 FR1;
- **Long training:** involved the full training described under phases III and IV.

**V. Seeking–taking–punishment phase:** Each cycle began exactly as described for the seeking–taking phase. However, during each punishment session, mild foot-shock punishment was delivered randomly on completion of 30% of the seeking cycles: the completion of the RI60 of 17 of the 25 total cycles resulted in the delivery of 0.1 mL of 15% EtOH following a taking lever response, whereas the completion of the RI60 of eight of the 25 cycles resulted in the random delivery of a 0.25–0.45 mA, 0.5 s foot shock. The taking lever was never presented if a seeking cycle resulted in punishment (so that punishment was never associated with the reinforcer). Although the cycles were randomly punished during a session, the first cycle of the session was always reinforced, and no more than two consecutive seeking cycles were punished. The intensity of the shock was progressively increased over daily sessions from 0.25 to 0.45 mA in 0.05 mA increments, before stabilizing at 0.45 mA for 6 consecutive daily sessions.

##### *Assessment of individual variability in resistance to outcome devaluation over the course of the development of compulsive alcohol seeking behaviour*

The training protocol was adapted from Olmstead et al.[6], Giuliano et al.[4,5], and Zapata et al.[7] and is illustrated in **Figure 1**.

After either **Short** or **Long training** under the seeking–taking task, two outcome devaluation assessments were conducted. Each assessment consisted of the following four steps[6,7]:

- **Devaluation of the outcome of drug seeking by extinction of drug taking**: To devalue the outcome of seeking responses, responding on the drug taking lever was extinguished. Rats underwent daily 1-h extinction sessions in which only the drug taking lever was present and lever responding had no programmed consequences (i.e. alcohol was not delivered). The total number of extinction sessions varied according to each subject's performance and ended when their lever presses had decreased by 90% from their performance on the first day of extinction of drug taking.
- **Test on the seeking lever**: Drug seeking was assessed in a 5-min test in which only the seeking lever was present under extinction conditions. If seeking behaviour is mediated by an action–outcome representation, seeking responses should decrease, because the outcome of seeking had been devalued by extinction of this outcome (presentation of the taking lever and access to alcohol)[8,9]. However, if seeking responses are unaffected or minimally affected by extinction of taking lever responses, this provides evidence that seeking behavior is mediated by a stimulus–response (habit) association[8-10].
- **Revaluation of the outcome of drug seeking**: The outcome of drug seeking was revalued by two sessions under FR1, where only the taking lever was present and each lever press resulted in 5-s CS illumination, extinction of the house-light, and delivery of 0.1 mL 15% EtOH. Rats were limited to a maximum of 45 rewards per 2h session.
- **Test on the seeking lever**: Drug seeking was again assessed in a 5-min test, in which only the drug seeking lever was present and responses did not deliver the taking lever or access to alcohol.

##### ***Assessment of alcohol drinking after development of compulsive alcohol seeking behaviour***

Once compulsive punishment-resistant alcohol seeking had developed, drinking behaviour was tested under different conditions:

- Rats underwent two sessions with 4h access to a 15% EtOH bottle either in the home-cage or in the operant chamber where they had been trained to seek alcohol. Alcohol intake (g/kg) was assessed for 4h. All rats were exposed to the two conditions in a counterbalanced order.
- Rats underwent either 5 or 15 cycles of the seeking–taking chained schedule under RI60/FR1. Once the cycles were completed, they remained in the operant chambers and had 4h access to a 15% EtOH bottle. All rats were exposed to the two conditions in a counterbalanced order. The tests sessions were preceded by three re-baseline sessions under RI60, and single re-baseline sessions were also interspersed between each test session.

##### *Assessment of quinine-resistant drinking after development of compulsive alcohol seeking behaviour*

After two additional re-baseline sessions of the seeking–taking chained schedule under RI60/FR1, compulsive drinking behaviour was tested under different conditions:

- Rats underwent 15 cycles of the seeking–taking chained schedule (RI60) followed by 30 min access to 0.1 g/L quinine-adulterated 15% EtOH in the same operant chamber in which the seeking–taking sessions occurred.
- Rats underwent 15 cycles of the seeking–taking chained schedule (RI60) in which the volume of quinine-adulterated alcohol delivered at the completion of each cycle was different from the baseline sessions. Taking lever responses resulted, via an FR1 schedule, in the illumination of the stimulus light above the taking lever for 5 s, delivery of 0.5 mL (rather than 0.1 mL as before) of 0.1 g/L quinine in 15% EtOH solution.

##### *Instrumental contingency degradation after development of compulsive alcohol seeking behaviour*

After four additional re-baseline sessions of the seeking–taking chained schedule under RI60/FR1, the fixed-ratio requirement of the taking response was increased to

3, and then to 5, in order to increase the effort required to obtain alcohol (0.1 mL 15% EtOH per cycle).

During the three subsequent sessions, in addition to alcohol being delivered as usual contingent on the taking response, additional volumes of 0.1 mL 15%EtOH were delivered noncontingently when the seeking lever was extended. Thus, 0.1 mL 15% EtOH was delivered after 5 taking responses on completion of each seeking–taking cycle, but the same volume was also noncontingently delivered under a random-ratio 60 s (RR60) schedule, independent of seeking lever responses. In this way, the seeking–taking link in the chain was degraded, enabling the assessment of the associative structure underlying seeking responses: action–outcome or stimulus–response[6-8,11,12]. The sessions lasted 1h, which was the average time rats took to complete 15 cycles under baseline conditions.

#### Data and Statistical analyses

Assumptions for normal distribution and homogeneity of variance were verified using the Shapiro–Wilk and Levene tests, respectively.

Two-step *k*-mean cluster analyses were performed to identify sub-populations of rats[3,4], based on the completed cycles over the last three days of exposure to punishment (0.45 mA), and to confirm the nature of the devaluation-sensitive and devaluation-resistant sub-populations identified after long training.

The effect of training (short vs long) on outcome devaluation was analysed by two-way repeated measures analyses of variance (ANOVA) with training (short vs long) and devaluation (revalued control vs devalued) as within-subject factors. The comparison of alcohol-seeking responses under devalued and revalued conditions was done using a paired *t* test carried out separately after short and long training. The effect of time on extinction of the taking responses was initially analysed by repeated measures ANOVA with time as a within-subject factor and a between-subjects factor of group (devaluation-sensitive (DS) vs resistant (DR) or non-compulsive (NC) vs compulsive (C)) was added to the model to test for possible differences between groups in extinction. The same strategy was adopted for the analysis of performance under the two-bottle choice procedure, seeking–taking, and seeking–taking–punishment chained procedures. A linear regression analysis was conducted to test whether alcohol intake prior to the establishment of compulsive seeking predicted later compulsive alcohol seeking. Pearson correlations were used to assess the relationship between the number of completed cycles on the last three days of 0.45 mA punishment and performance on the first day of extinction.

When the data were expressed as percentage change from baseline, one-way ANOVA was carried out with group (devaluation-sensitive (DS) vs -resistant (DR) and non-compulsive (NC) vs compulsive (C)) as a between-subjects factor. The effect of quinine adulteration was analysed by two-way ANOVA with time (early vs late), context (home cage vs operant chamber), or cycle (5 vs 15) as a within-subject factor and group (C vs NC rats) as a between-subjects factor. The effect of contingency degradation on seeking and taking responses (expressed as percentage change from baseline), and on the number of contingent and non-contingent reinforcers obtained, were analysed by two-way ANOVA with time (3 sessions) as a within-subject factor and group (C vs NC rats) as a between-subjects factor.

### Results

#### *Training to develop compulsive alcohol seeking.*

##### *I.*Two-bottle choice procedure

Data from the two-bottle choice procedure, and from the instrumental training (leading to the assessment of the compulsive nature of alcohol seeking represented in the Figures), were systematically analysed: (i) considering the "whole-group", n=25; (ii) retrospectively considering the "sub-populations" identified according to resistance to outcome devaluation after long training, i.e. devaluation sensitive (DS, n=8) vs devaluation resistant (DR, n=17); (iii) retrospectively considering the sub-populations identified according to punishment-resistant seeking behaviour, i.e. compulsive (C, n=7) vs non-compulsive (NC, n=10).

Subjects underwent an intermittent two-bottle choice procedure for 12 sessions over which alcohol intake (measured every 24h) progressively increased across the various groups **(Figure S1A)** ["**whole group"**: time: $F_{11,264} = 10.40$, $p< .001$, $\eta_p^2= .30$; **DS vs DR**: time: $F_{11,253} = 14.60$, $p< .001$, $\eta_p^2= .39$; group: $F_{1,23} < 1$, $\eta_p^2= .001$; time x group: $F_{11,253} = 3.83$, $p< .001$, $\eta_p^2= .14$; **NC vs C**: time: $F_{11,165} = 7.01$, $p< .001$, $\eta_p^2= .32$; group: $F_{1,15} < 1$, $\eta_p^2= .06$; time x group: $F_{11,165} < 1$, $\eta_p^2= .06$]. As compared to that shown on day 1, daily alcohol intake increased to a stable level from day 3 or day 6 for the entire population ($p< .021$), DS ($p< .018$) and NC rats ($p< .019$), respectively. DR and C rats already showed a high level of drinking on the first session that did not differ from that seen during subsequent sessions. Additionally, during the first session rats later showing higher sensitivity to outcome devaluation (DS) had a lower alcohol intake than their counterparts ($p= .008$), but it quickly leveled out.

Accordingly, preference for alcohol over water increased over time across groups **(Figure S1B)** [**"whole-group"**: time: $F_{11,264} = 23.89$, $p< .001$, $\eta_p^2= .50$; **DS vs DR:** group: $F_{1,23} = 1.11$, NS; block: $F_{3,69} = 17.77$, $p< .001$, $\eta_p^2= .44$; block x group: $F_{3,69} = 4.90$, $p= .004$, $\eta_p^2= .018$; **NC vs C:** group: $F_{1,15} = 2.13$, $p= .16$, $\eta_p^2= .12$; block: $F_{3,45} = 8.76$, $p< .001$, $\eta_p^2= .37$; block x group: $F_{3,45} = 1.30$, NS]. Hence, alcohol preference was stable and higher than on the first day from day 3 onwards in the entire group as well as each subpopulation, save that the increased preference in DR rats was evident from day 6 onwards. As for alcohol intake, C rats had a preference for alcohol from day 1 that did not differ from that shown during the next 11 sessions, and DS rats showed initially less preference for alcohol than DR rats on the first session [$p= .009$].

##### *II.*Seeking–Taking chained schedule

Rats were trained instrumentally to respond for alcohol under continuous reinforcement[13] for four sessions. By the end of this, they all obtained the maximum number of rewards available per session (i.e. 45 reinforcers) **(Figure S2A)** [main effect of time: $F_{3,66} = 7.82$, $p= .001$, $\eta_p^2= .026$, showing full acquisition at day 3 as revealed by a post-hoc analysis with days 3 and 4 compared to day 1: $p= .003$, and $p= .002$, respectively]. They then underwent three 2-hour sessions under a seeking–taking chained schedule of reinforcement (RI5/FR1)[4,5,14], eventually reaching $113 \pm 15.89$ and $35 \pm 4.22$ lever presses on the seeking and taking levers, respectively **(Figure S2B)**. Withholding alcohol delivery resulted in the progressive extinction of the responses on the taking lever[6,7] over 17 sessions [time: $F_{16,384} = 86.01$, $p< .001$, $\eta_p^2= .78$ and time: $F_{16,384} = 138.32$, $p< .001$, $\eta_p^2= .85$, for taking responses and %change from day 1, respectively]. Rats had decreased their response level by ~50% by the second session [vs day1: $p< .001$ and $p< .001$, for taking responses and %change from day 1 respectively] which further decreased in subsequent sessions. By the last two extinction sessions, responding had decreased by more than 90%, with subjects making on average 9 ± .97 taking responses **(Figure S2C-D)**.

After the first devaluation test, training under the seeking–taking chained schedule of reinforcement resumed and reached the final stage of RI60/FR1, under which rats made 320 ± 35.07 and 24 ± .34 seeking and taking responses per session, respectively, as measured over the last two sessions of training (in which the number of cycles was limited to 25) **(Figure S2E)**. Withholding alcohol delivery again resulted in the progressive extinction of taking responses over 24 sessions [time: $F_{23,552} = 91.82$, $p< .001$, $\eta_p^2= .79$ and time: $F_{23,552} = 121.84$, $p< .001$, $\eta_p^2= .83$, for taking responses and %change from day 1, respectively]. On the second and third extinction session rats decreased their responding to 40% and 60% of that seen on the first day [vs day1: $p< .001$ and $p< .001$, for taking responses and %change from day 1, respectively], eventually decreasing by more than 90% to make 10 ± .98 taking responses during the last two sessions **(Figure S2F-G)**.

Rats that were subsequently identified as being sensitive or resistant to outcome devaluation did not differ in extinction of their taking responses during either early **(Figure S3A-B)** or late extinction training **(Figure S3C-D)** [**EARLY, after short training**: time: $F_{16,368} = 69.53$, $p< .001$, $\eta_p^2= .75$; group: $F_{1,23} < 1$, $\eta_p^2= .002$; time x group: $F_{16,368} < 1$, NS and time: $F_{16,368} = 115.47$, $p< .001$, $\eta_p^2= .83$; group: $F_{1,23} < 1$, NS; time x group: $F_{16,368} < 1$, NS for taking responses and %change from day 1, respectively; **LATE, after long training**: time: $F_{23,529} = 78.80$, $p< .001$, $\eta_p^2= .77$; group: $F_{1,23} < 1$, NS; time x group: $F_{23,529} < 1$, NS and time: $F_{23,529} = 106.94$, $p< .001$, $\eta_p^2= .82$; group: $F_{1,23} < 1$, NS; time x group: $F_{23,529} = 1.23$, NS for taking responses and %change from day 1, respectively]. Nor did they differ in their acquisition of alcohol self-administration **(Figure S3E)** or performance under the seeking taking task [**short training**, seeking responses: group: $F_{1,23} < 1$, NS; taking responses: group: $F_{1,23} < 1$, NS **(Figure S3F)**; **long training**, seeking responses: group: $F_{1,23} = 2.30$, NS; taking responses: group: $F_{1,23} = 1.59$, NS **(Figure S3G)**].

In marked contrast, rats that subsequently developed compulsive alcohol seeking behaviour and their non-compulsive counterparts differed in their response to the extinction of the taking lever after both short- [time: $F_{16,240} = 55.20$, $p< .001$, $\eta_p^2= .79$; group: $F_{1,15} < 1$, NS; time x group: $F_{16,240} = 2.89$, $p= .02$, $\eta_p^2= .16$ and time: $F_{16,240} =$

80.41, $p< .001$, $\eta_p^2= .84$; group: $F_{1,15} = 3.99$, $p= .064$, $\eta_p^2= .21$; time x group: $F_{16,240} = 1.12$, NS for taking responses and %change from day 1, respectively] **(Figure S4AB)** and long-term training [time: $F_{23,345} = 65.02$, $p< .001$, $\eta_p^2= .81$; group: $F_{1,15} = 1.13$, NS; time x group: $F_{23,345} = 2.67$, $p= .03$, $\eta_p^2= .15$ and time: $F_{23,345} = 73.91$, $p< .001$, $\eta_p^2= .83$; group: $F_{1,15} = 1.16$, NS; time x group: $F_{23,345} < 1$, NS for taking responses and %change from day 1, respectively] **(Figure S4CD)**. In both instances this was due to C rats persisting in responding more on the first extinction session than NC rats [$p= .034$ and $p= .075$, respectively], also supported by the fact that responding by NC rats significantly differed from day 1 in each subsequent extinction session [all $p< .028$] whereas it only decreased significantly in C rats from the third session onwards [all $p< .0001$]. This response stickiness shown by C rats was not related to a different ability to learn the new response non-US association that underlies extinction, as both groups then exhibited a similar pattern of decease in their response level across daily sessions, eventually reaching a 90% decrease by the last sessions. Nor was it due to a difference in the acquisition of alcohol self-administration **(Figure S3E)** or performance in the seeking–taking task [**short training,** seeking responses: group: $F_{1,15} = 1.72$, NS; taking responses: group: $F_{1,15} < 1$, NS, **Figure S3F)**; **long training,** seeking responses: group: $F_{1,15} < 1$, NS; taking responses: group: $F_{1,15} = 2.19$, NS **(Figure S3G)**].

##### *III.* Identification of sub-populations according to punishment-resistant alcohol seeking

After completing the second devaluation test, rats resumed the instrumental training so the compulsive nature of their alcohol seeking behaviour could be measured. Once punishment was introduced, daily sessions were limited to a maximum of 25 cycles, of which 17 were reinforced and 8 were randomly punished. The completion of the 25 cycles/session was an index of animals' persistent alcohol seeking in the face of negative consequences, i.e. the degree to which they took the risk of being unpredictably punished in order to take and drink alcohol[4,5,15]. Over the course of the punished sessions, with shock intensity increasing from 0.25 to 0.45 mA between sessions, rats progressively decreased the number of completed cycles [time: $F_{10,150} = 17.62$, $p< .001$, $\eta_p^2= .54$], which were significantly less than those obtained under baseline conditions by session 5 (the first day at 0.45 mA) onwards [$p< .005$].

A cluster analysis, performed on the number of cycles completed during the last three sessions of the seeking–taking–punishment task, segregated three subgroups of rats: 7 rats displayed alcohol seeking that was completely resistant to punishment, i.e., was compulsive (C), while 10 rats showed a marked decrease in alcohol seeking and were therefore considered non-compulsive (NC). The remaining 8 rats responded under punishment at intermediate levels (I). As expected by the definition of compulsive versus non-compulsive responding, C and NC rats (only these two subgroups were considered for all the between-subject comparisons in the present study) showed a very different behavioral response to punishment [time: $F_{9,135} = 14.26$, $p < .001$, $\eta_p^2 = .49$; group: $F_{1,15} = 118.74$, $p < .001$, $\eta_p^2 = .89$; time x group: $F_{9,135} = 7.93$, $p < .001$, $\eta_p^2 = .35$], with NC rats greatly decreasing the number of completed seeking–taking cycles in the face of punishment and C rats maintaining a constant level of completed cycles throughout **(Figure S5A)**. This differential decrease in completed seeking–taking cycles reflected a differential vulnerability to persist in seeking alcohol even though this incurred the receipt of shocks [time: $F_{9,135} = 8.71$, $p < .001$, $\eta_p^2 = .37$; group: $F_{1,15} = 16.66$, $p = .001$, $\eta_p^2 = .53$; time x group: $F_{9,135} = 2.04$, $p = .039$, $\eta_p^2 = .12$] **(Figure S5B),** resulting in a lower number of opportunities to take alcohol [time: $F_{9,135} = 12.66$, $p < .001$, $\eta_p^2 = .46$; group: $F_{1,15} = 112.99$, $p < .001$, $\eta_p^2 = .88$; time x group: $F_{9,135} = 7.39$, $p < .001$, $\eta_p^2 = .33$] **(Figure S5C)** and receive shocks [time: $F_{9,135} = 11.91$, $p < .001$, $\eta_p^2 = .44$; group: $F_{1,15} = 116.31$, $p < .001$, $\eta_p^2 = .89$; time x group: $F_{9,135} = 5.67$, $p < .001$, $\eta_p^2 = .27$] **(Figure S5D)**. The differential response to probabilistic punishment of NC and C rats was, however, not due to a difference in baseline seeking performance, as their seeking [group: $F_{1,15} = 2.19$, NS] and taking responses [group: $F_{1,15} < 1$, NS] before punishment exposure were not significantly different.

#### Supporting figures

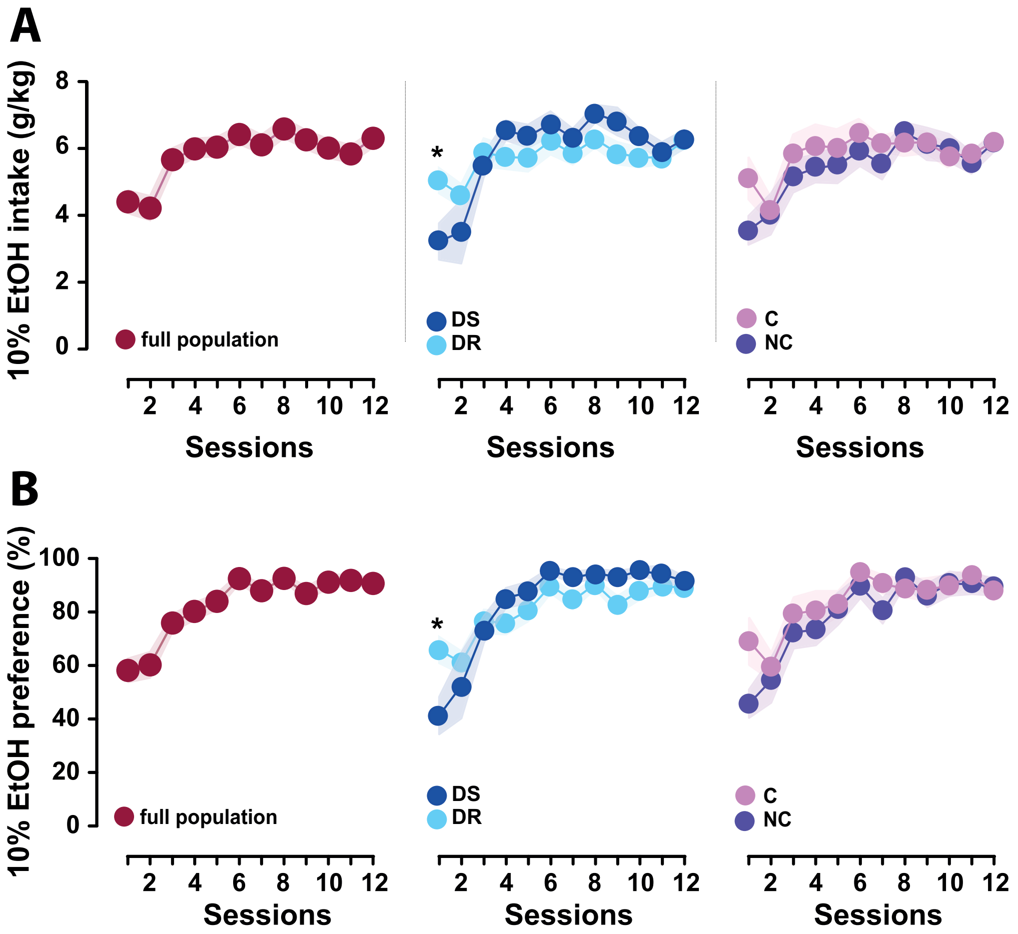


**Figure S1: Acquisition of alcohol drinking in a two-bottle choice procedure across groups.**

The alcohol-preferring phenotype of the 25 rats was assessed under the two-bottle choice procedure. During the 12 24-h sessions of intermittent access to EtOH and water, all the rats reached similar levels of alcohol intake and preference for EtOH over water. Alcohol intake (g/kg) **(A)** and EtOH preference (%, w/w) **(B)** increased over 12 sessions at the whole-group level (n=25, in burgundy, **left panel**), as well as when subpopulations were considered independently, namely with respect to their sensitivity to outcome devaluation (**middle panel**) or to punishment (**right panel**). Individuals later identified in outcome devaluation tests as devaluation sensitive (DS, n=8, in blue), or devaluation resistant (DR, n=17, in light blue) showed a difference in their initial level of alcohol intake and preference [*: vs DS, p< .01]. No differences were observed between non-compulsive (NC, n=10, in purple) and compulsive rats (C, n=7, in pink). Graphs show means ± SEM.

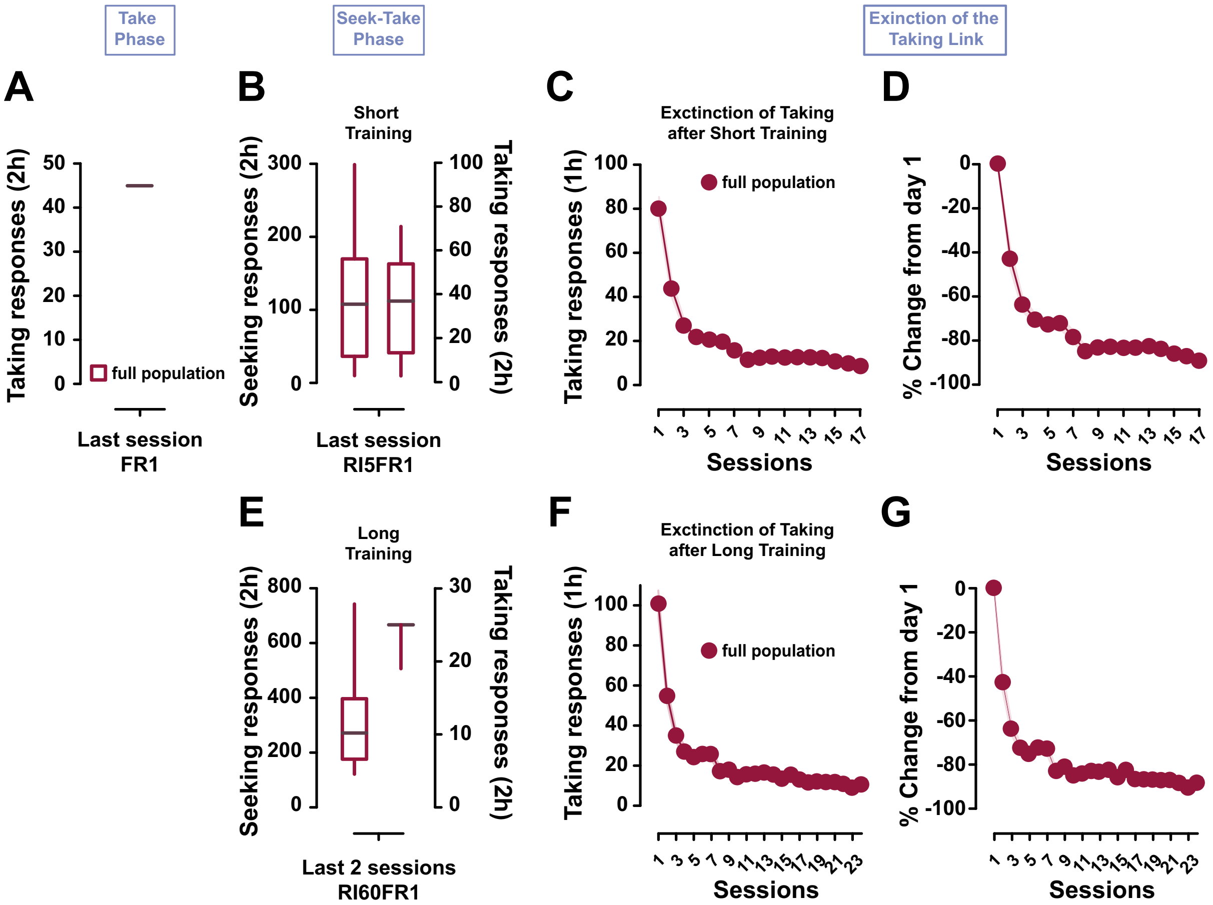


**Figure S2: Whole-group analysis of performance during instrumental training and extinction of the taking lever.**

Alcohol-preferring (P) rats (n=25) were trained under FR1 ("Take Phase") **(A)**. They were then trained under a seeking–taking chained schedule of reinforcement (RI5/FR1) for three sessions ("Short Training"). Seeking (left) and taking (right) responses are shown for the last session **(B)**. The taking link was then extinguished (i.e. no alcohol was delivered) over 17 sessions before the first devaluation test was carried out. Taking responses **(C)** and percentage change from the first day of extinction **(D)** are shown. The instrumental training under the seeking–taking schedule resumed, to reach the final stage under RI60/FR1 ("Long Training"). Seeking (left) and taking (right) responses are shown for the last session **(E)**. The taking link was then extinguished (i.e. no alcohol was delivered) over 24 sessions before the first devaluation test was carried out. Taking responses **(F)** and percentage change **(G)** are shown for the first day of extinction. Graphs show means ± SEM and box plots show the median, minimum, maximum, and quartiles.

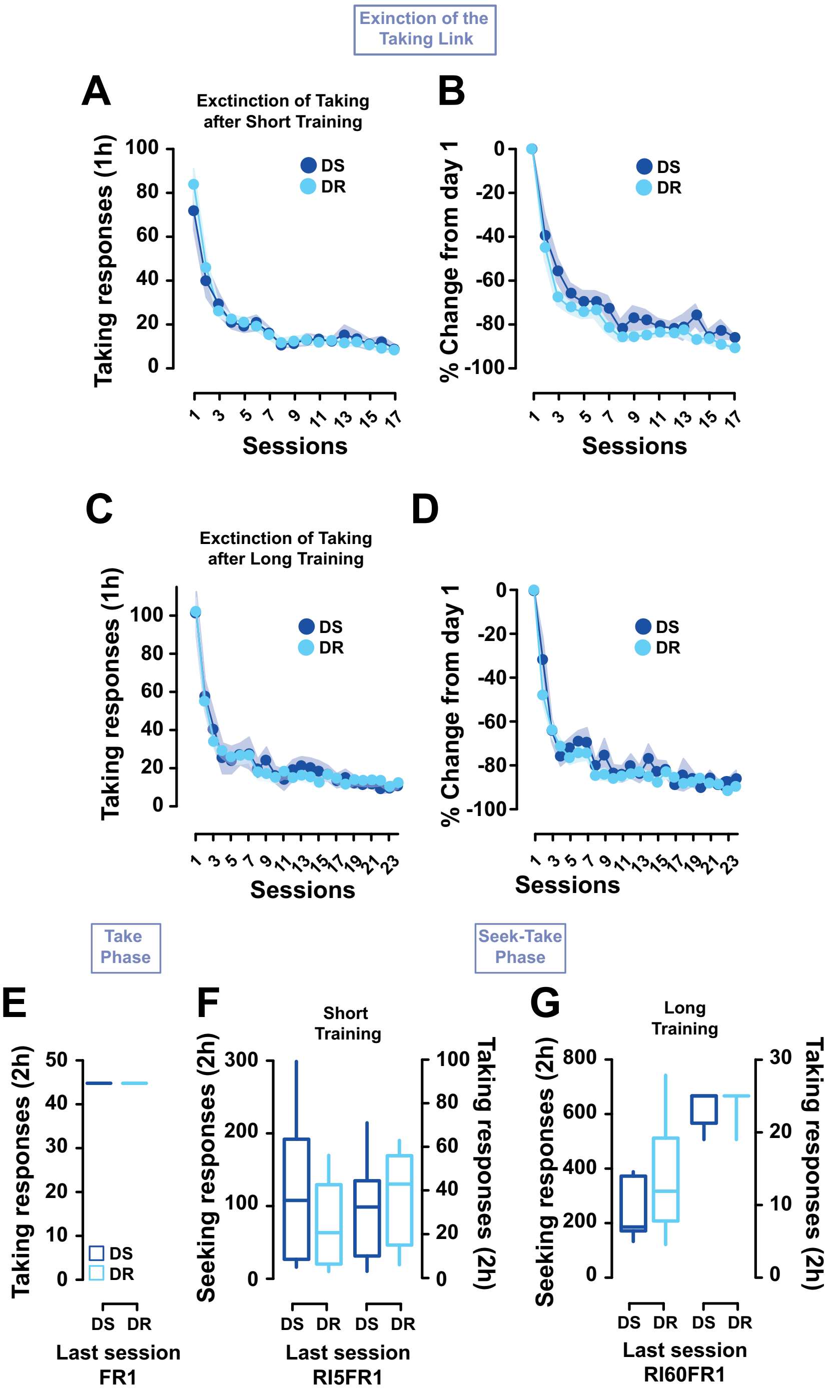


**Figure S3: Devaluation-sensitive and devaluation-resistant rats did not differ in their extinction of alcohol taking responding and they acquired alcohol self-administration and alcohol seeking similarly.**

Devaluation-sensitive (DS, blue, n=8) and devaluation-resistant (DR, light blue, n=17) rats extinguished their taking responses similarly over the 17 and 24 extinction sessions that were used after short-term **(A-B)** and long-term training **(C-D)**, respectively to devalue the outcome of the seeking response (namely the taking lever).

Similarly, DS and DR rats did not differ in their acquisition of alcohol self-administration, shown by the maximum number of rewards acquired during the last session under FR1 **(E)**, or in their performance under the seeking–taking task, either after short training **(F)** or long training **(G)** (plots show performance at the end of training). Graphs show means ± SEM and box plots are as before.

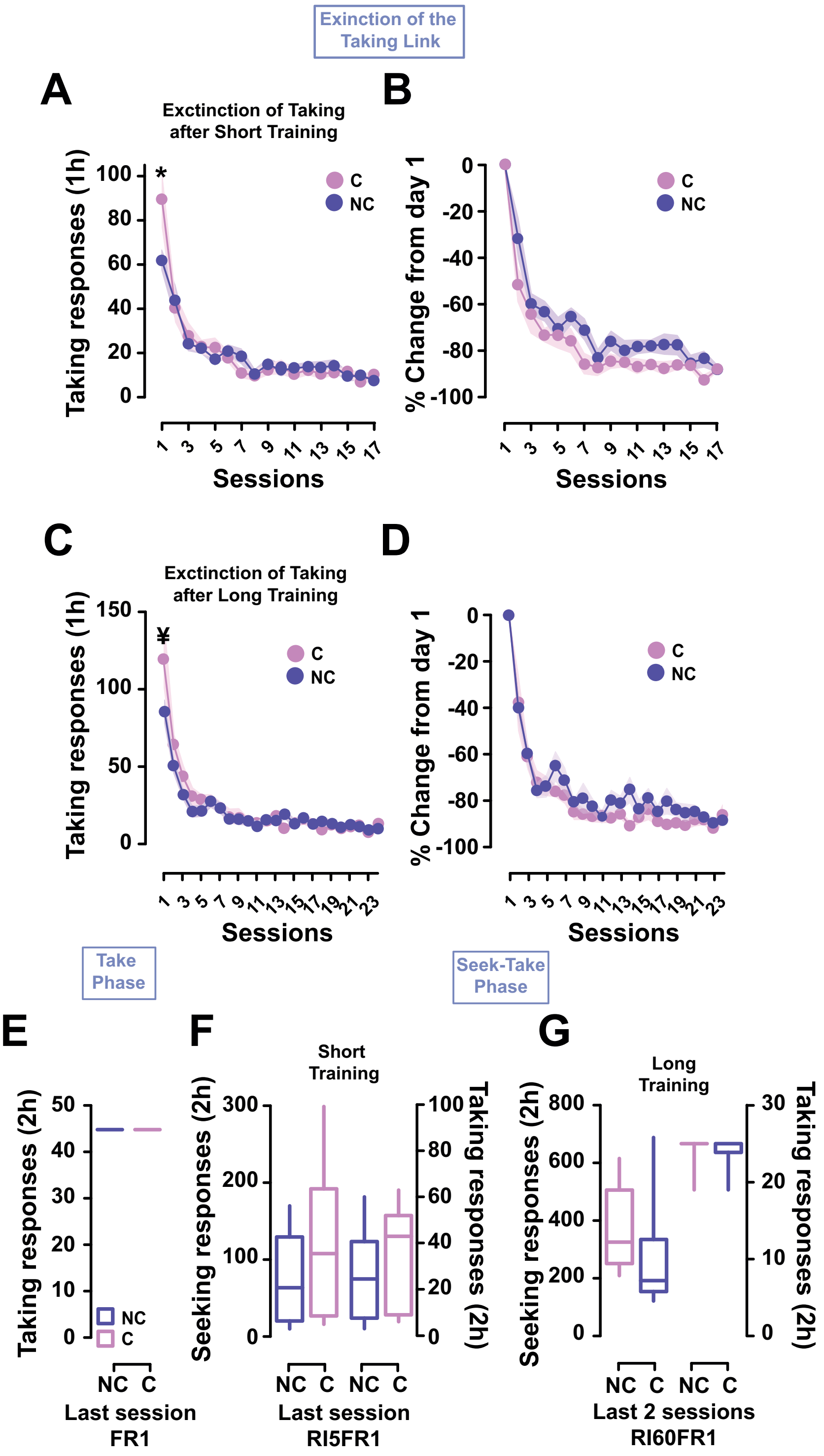


**Figure S4: Compulsive and non-compulsive rats did not differ in their seeking or taking responses during training, but differed slightly in their propensity to extinguish alcohol taking.**

Compulsive (C, pink, n=7) and non-compulsive (NC, purple, n=10) subjects displayed a different response to the extinction of the taking lever over the 17 and 24 extinction sessions that were used after short-term **(A-B)** and long-term training **(C-D)**, respectively. C rats responded more during the first session of extinction training [*: vs NC, p= .034 and ¥: vs NC, p= .075].

However, NC and C rats did not differ in their ability to extinguish subsequently, and they had both decreased their response rate by 90% towards the end of the extinction periods **(B** and **D)**.

NC and C rats did not differ in their acquisition of alcohol self-administration (number of rewards obtained during the last session under FR1; **E**) or in their performance under the seeking–taking task, either after short training **(F)** or long training **(G)** (performance shown at the end of training). Graphs show means ± SEM and box plots.

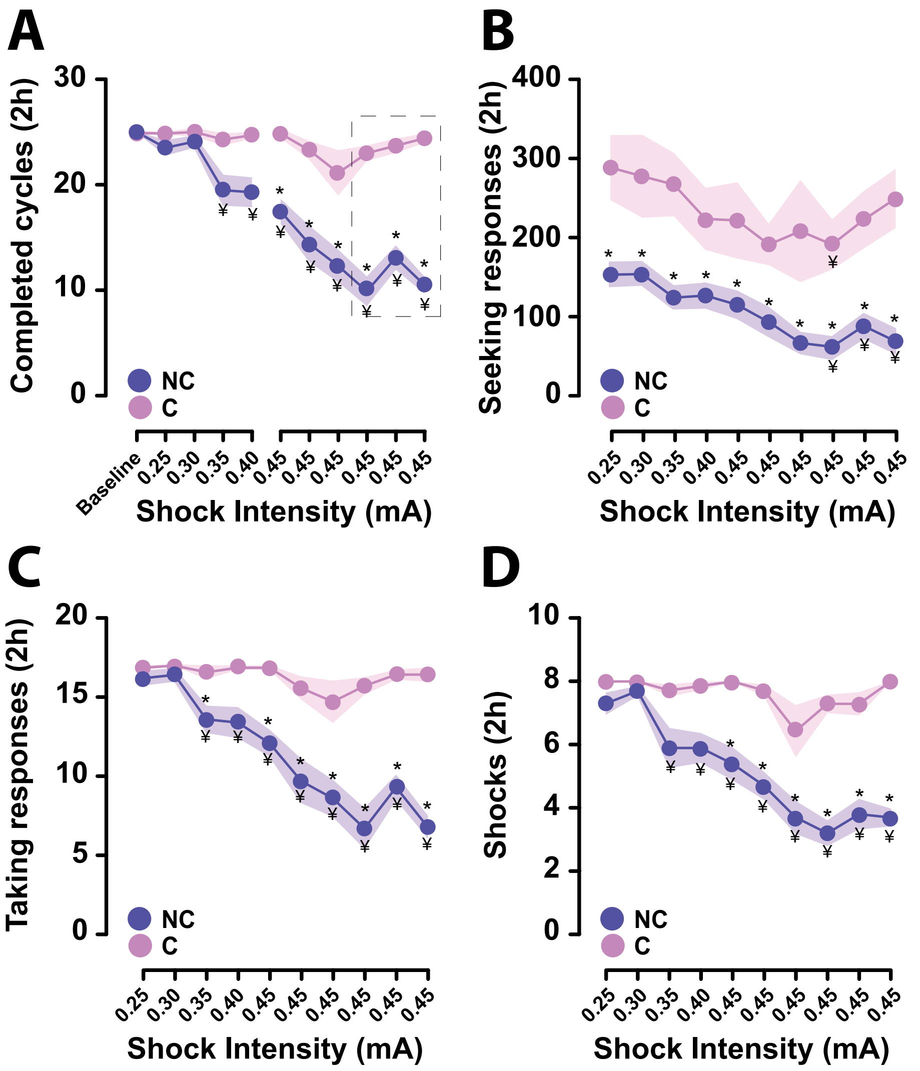


**Figure S5: Identification of compulsive and non-compulsive individuals**

P rats (n=25) seeking alcohol under a seeking–taking schedule were challenged to persist in responding on the seeking lever despite the risk of punishment by mild electric foot-shocks. Shocks were delivered under a probabilistic schedule, with intensity increasing over daily sessions from 0.25 mA, via 0.30, 0.35, and 0.40 mA, and eventually to 0.45 mA for 6 consecutive sessions. Based on the persistence of alcohol seeking during the last three punishment sessions (measured by the number of completed seeking–taking cycles), a cluster analysis enabled the segregation of rats into three subgroups, of which the compulsive (C, pink, n=7) and non-compulsive (NC, purple, n=10) are compared here. **(A)** The differential resistance to punishment emerged on the third punishment session (0.35 mA) from which point NC rats differed from C rats [¥: vs NC, $p < .01$]. C rats maintained responding at a level similar to their baseline, while NC rats had decreased responding by the first 0.45 mA punishment session [*: vs baseline, $p \leq .02$]. This decrease in the total number of completed cycles was reflected in a decrease, for NC rats, in seeking [*: vs day 1, $p \leq .036$ and ¥: vs NC, $p \leq .03$] **(B)** and taking responses [*: vs day 1, $p \leq .026$ and ¥: vs NC, $p \leq .014$] **(C)**, with a corresponding decrease in the number of shocks received overall [*: vs day 1, $p \leq .04$ and ¥: vs NC, $p \leq .03$] **(D).** Graphs show means ± SEM.
